# Comparative genome analysis using sample-specific string detection in accurate long reads

**DOI:** 10.1101/2021.03.23.436571

**Authors:** Parsoa Khorsand, Luca Denti, Human Genome Structural Variant Consortium, Paola Bonizzoni, Rayan Chikhi, Fereydoun Hormozdiari

**Affiliations:** Genome Center, UC Davis, Davis, CA, USA; Department of Computational Biology, Institut Pasteur, Paris, France; Department of Informatics, Systems and Communication, University of Milano-Bicocca, Milano, 20126, Italy; UC Davis MIND Institute, Sacramento, CA, USA; Department of Biochemistry and Molecular Medicine, Sacramento, UC Davis, Sacramento, CA, USA

## Abstract

**Motivation:** Comparative genome analysis of two or more whole-genome sequenced (WGS) samples is at the core of most applications in genomics. These include discovery of genomic differences segregating in population, case-control analysis in common diseases, and rare disorders. With the current progress of accurate long-read sequencing technologies (e.g., circular consensus sequencing from PacBio sequencers) we can dive into studying repeat regions of genome (e.g., segmental duplications) and hard-to-detect variants (e.g., complex structural variants).

**Results:** We propose a novel framework for addressing the comparative genome analysis by discovery of strings that are specific to one genome (“samples-specific” strings). We have developed an accurate and efficient novel method for discovery of samples-specific strings between two groups of WGS samples. The proposed approach will give us the ability to perform comparative genome analysis without the need to map the reads and is not hindered by shortcomings of the reference genome. We show that the proposed approach is capable of accurately finding samples-specific strings representing nearly all variation (*>* 98%) reported across pairs or trios of WGS samples using accurate long reads (e.g., PacBio HiFi data).

**Availability:** The proposed tool is publicly available at https://github.com/Parsoa/PingPong.

## 1 Introduction

Whole-genome sequencing (WGS) has become the dominant approach in studying variations across genomes. Today, WGS data continues to provide invaluable insight into every aspect of biology. In particular, comparative analysis of multiple samples using WGS data is fundamental in understanding genetics of disorders, traits, and evolution. The comparison of differences found between exome and genome of affected cases and unaffected controls has successfully found genetic variants associated with disorders and guided predicting genes contributing to disorders [12]. Population genomics studies benefit from WGS data by finding sequences and variants shared or discriminative between different populations [13, 30]. Furthermore, evolutionary studies also benefit from such comparative studies in a multiple species setting [41, 35].

High-throughput short-read sequencing (i.e., Illumina) has been the driving force behind most of the WGS studies in the past decade. Short-read sequencing is cheap, provides high-throughput data, and has low error rate [45]. However, it also has several major drawbacks. First, the assembly of the eukaryotic genomes using short-read sequencing data is non-trivial and computationally resource intensive [21]. Second, the short length of the reads (generally below 250bp) produced by these technologies has caused significant complexity and ambiguity in studying repeat regions of the genome [29, 51, 9]. Third, the quality of structural variation (SV) and other complex variant calls predicted using short-reads data has remained low despite significant bioinformatics efforts, and requires orthogonal validations [46, 10]. Finally, several types of genetic variations are hard to predict using short-read sequencing technologies due to their repeat nature (e.g., VNTR expansions [3]).

With the advent of long-read sequencing technologies (e.g., PacBio or Oxford Nanopore) we have access to long reads (*>* 10 kbp) that can be used to overcome the above-mentioned shortcomings of short-read sequencing [32, 10, 7]. WGS data from long-read sequencing technologies enables one to discover and further study variants that were either hidden or unreliably predicted from short-read data. For example, a recent benchmark showed that long-read sequencing data enabled to find a large fraction (over 50% of SVs) which were unreported from short-read sequencing data [10, 9, 51].

One of the main objectives of performing WGS is the comparison of two or more genomes. Such comparative genomics studies are concerned with multiple individuals from the same species or from multiple species, either in a case versus control setting or within population genomics [18]. Discovery of differences between multiple samples using WGS is at the core of most genomic analysis.

The dominant framework for comparative variant analysis among multiple sequenced samples is based on mapping the reads to the reference genome [24, 28], predicting variants in each sample, and comparing the predicted loci [39, 9, 31, 1]. The comparison of variants predicted between multiple samples is based on overlapping the predicted variant locations. This strategy is effective for comparing SNVs, however for many SVs the exact breakpoint position is hard to establish and ambiguities can negatively affect accuracy. There are several heuristics used for comparing SVs in multiple samples by considering that the exact breakpoint for the SV might not be known or ambiguous^1^. They are based on merging SVs with approximately close breakpoints and considering reciprocal overlaps as a match [10]. These heuristics tend to work for SVs in simple regions of the genome. However, for more complex scenarios such as STR/VNTR expansions [17, 4], SVs with adjacent SNP variants [8], or SVs with breakpoints in repeats (e.g., segmental duplications) will result in reduction of accuracy as these heuristics tend to fail [10, 46, 34].

An alternative approach for comparative genome analysis is not to compare the predicted variants among multiple samples but to find the **sequences containing breakpoints** that are different between samples. This approach can be done without the need to map the reads to the reference genome and predicting variants per sample (i.e., mapping-free approaches). Examples of mapping-free approaches for studying genomes are DiscoSNP++ [37], Scalpel [33], LAVA [44], VarGeno [47], MALVA [14], Nebula [19] and HAWK [43] for detection and genotyping of variants in WGS data, and DE-Kupl [2] for detecting RNA-Seq variations. These mapping-free approaches have the advantage of not being impacted by the possibility of ambiguity in SV breakpoints or inaccuracies in the reference genome itself. The mapping-free approaches developed for studying short-read sequencing data are mostly based on finding *k*-mers that distinguish one sample from other samples. The idea of computing k-mers that are unique to a target w.r.t. a background set of genomes is also proposed in [38]. In general, the length of *k*-mers (i.e., *k*) is a fixed constant and usually short. However, for long and accurate reads we are not limited by the length of the short reads and can select arbitrarily long *k*-mers if needed. This flexibility on length of sequences selected can be advantageous for comparative studying of repeat regions of the genome. The tools mentioned above are fundamentally unable to deal with variable-length *k*-mers and therefore novel developments are needed to fully explore this direction.

We propose a novel method for comparative analysis of multiple WGS samples using accurate long-read sequencing data (e.g., HiFi reads from PacBio [49]), without the need to map the reads to a reference genome or choose a fixed *k* value. The advantages of utilizing flexible length strings (e.g., adaptive seeds) in pattern matching has been previously demonstrated [20]. The main novelty is the formulation and the resolution of a new computational problem concerned with enumerating sample-specific strings, while avoiding a combinatorial explosion due to the quadratic size of the set of potential candidates. We show that this approach enables identifying nearly all sequences spanning variants between two human genomes on actual PacBio HiFi data. Some of the applications of the proposed comparative genome analysis framework include finding *de novo* variants, sequences segregating in a pedigree, or markers distinguishing between populations (e.g., cases and controls).

## 2 Problem definition

Consider two sets of strings: *T* (targets) and *R* (references). Here by *references* we mean either 1) a reference genome, or 2) a set of unassembled reads that are coming from an unknown reference genome, or 3) a heterogeneous set of reads and genomes that are taken together to be the reference pangenome of some population of interest. We are interested in enumerating substrings of the targets that do not appear as exact substrings of the references.

As a motivating example consider two individuals, Tia and Red, and their respective sets of sequencing reads *T* and *R*. We define a **variant** as a genomic event that can be described by a single line in the VCF format, such as a single nucleotide polymorphism (SNP), an insertion or deletion, or a structural variant such as a duplication, or a translocation. More complex forms of genomic variation, e.g., an inversion-duplication, can be seen as combinations of variants and therefore are not further considered here. The intuition is that for each variant, there should exist at least one substring of the genome of Tia spanning this variant that is not found within the genome of Red. Indeed, the whole genome of Tia would be one such substring, but there also likely exist shorter strings than that. Translating this observation to reads, there should exist for each variant at least one substring of *T* that is not found in *R*. We postulate, and will later experimentally verify, that with long and accurate enough reads virtually all variants can be found in substrings of *T* that do not appear in *R*.

We are now returning to the abstract formulation of our initial problem of finding substrings of the targets not found in the references. For two strings *s* and *t*, we will use the notation *s* ⊏ *t* to indicate that *s* is a substring of *t* (and *s* 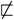 *t* for *s* is not a substring of *t*). Formally we want to enumerate the set 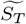 of all strings *s* such that

1. there exists *t* ∈ *T* where *s* ⊏ *t*, and
2. for all *r* ∈ *R*, 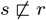.

In the worst case, the size of 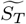 can be quadratic in the total length of strings in *T*, which is too large to be stored or even enumerated. Therefore we will instead seek a reduced set of strings *S*_*T*_ that can be seen as a minimal representation of 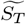 that do not consider strings having proper substrings in 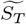 (the **substring-free** property). This is formalized as the following problem:

Problem 1 (Substring-Free Sample-specific (SFS) strings). Let *T* and *R* be two sets of strings, targets and references, respectively. *Let* 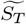 *be the set of all strings satisfying conditions 1 and 2 above. Return the largest subset* 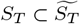 *such that for all s* ∈ *S*_*T*_, *there does not exist* 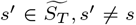, *where s* ′ *⊏ s; i*.*e*., *S*_*T*_ *is the set of all strings from* 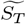 *for which no shorter string of* 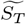 *is substring of them*.

A string *s* ∈ *S*_*T*_ is then called a *T-specific string* w.r.t. references *R*, or simply *specific string* when *T* and *R* are clear from the context. Furthermore, a *T*-specific string *s* that is a substring of *t* ∈ *T*, will be also called a *t*-specific string. In the following, we will sometimes omit recalling that *T*-specific (and *t*-specific) strings are substring-free.

We will refer to Problem 1 as the “SFS problem”, and an instance is illustrated in Figure 1a. It is easy to see that SFS can be (inefficiently) solved in *O*(*n*^3^) worst-case time and *O*(*m*^3^) memory, where *n* and *m* are the total lengths of strings in *T* and *R* respectively. The set 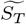 can be constructed by enumerating all substrings of *T* and checking their membership in a hash table containing all substrings in *R*; then another pass over 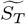 constructs *S*_*T*_ in linear time and space over the total length of strings in 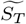, e.g., through indexing 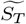 using a FM-index. In this paper, we will propose a novel and more efficient quadratic-time *O*(*n*^2^) algorithm (Algorithm 1 in Section 4) using linear-space *O*(*m*) for solving the SFS problem. We will also propose a heuristic version of the algorithm that solves a relaxed variant of Problem 1 in linear-time *O*(*n*). All these complexities are on top of the FMD-index construction [25], which in our case can be done in *O*(*m*) time and space [5].

**Fig 1(a):**
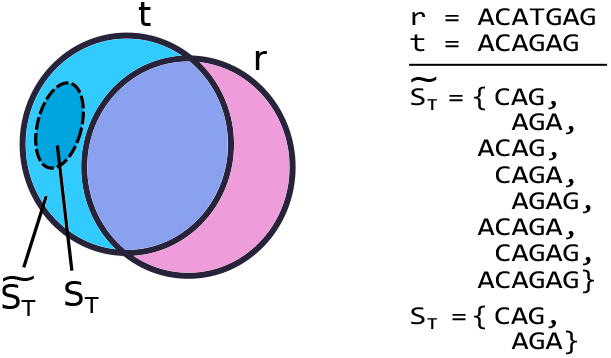
Illustration of the SFS framework. Consider a target string *t* and a reference string *r*, each represented by a circle symbolizing all substrings. Blue area: substrings of *t* not in *r*; pink: substrings of *r* not in *t*; purple: substrings common to both *t* and *r*. We start by enumerating 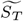, consisting of all strings *s* that satisfy conditions 1 and 2 of Section 2 (i.e., *s* is a substring of *t* and not a substring of *r*). Then, the set *S*_*T*_ (result of SFS) is the largest substring-free subset of 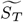.

**Fig 1(b):**
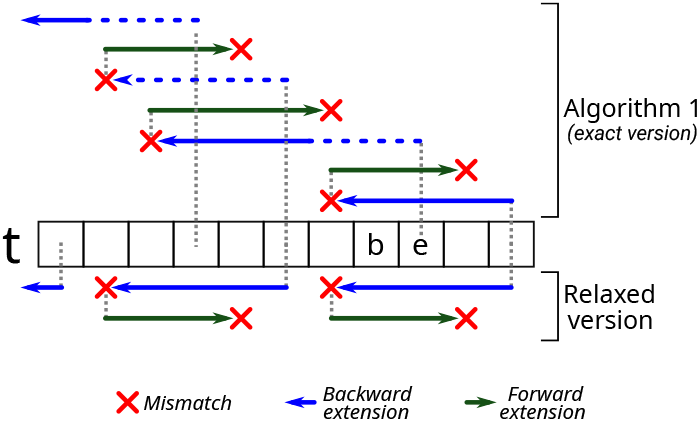
The Ping-pong search algorithm. (top side) starts from the end of the input string *t* and alternates between backward and forward extensions. When the backward extension (blue arrows) ends due to a mismatch (red cross), the algorithm starts a forward extension (green arrows) until another mismatch is found. After a single iteration (outer `while` loop of the pseudocode), a *t*-specific string *t*[*b* − 1 : *e* + 1] is found and the algorithm restarts the search from position *e*, allowing solutions to “overlap” on *t*. A dashed blue line represents bi-intervals that were already computed during a forward search (and therefore not recomputed in the next iteration). In the relaxed version of the algorithm (bottom side), solutions cannot overlap and the search restarts from position *b* − 2 instead of *e*. We note that Algorithm 1 outputs substring *t*[*b* : *e*] since *b* (resp. *e*) has been already previously decremented (resp. incremented).

The following property shows that it is sufficient to consider instances of the SFS problem where *T* is reduced to a single string.

Property 1 (Local substring-free property). *Let T and R be two sets of strings (targets and references, respectively). The set S*_*T*_ *of T-specific strings w*.*r*.*t. R, i*.*e*., *the solution of SFS problem, can be computed as the union of the sets S*_*t*_ *with t* ∈ *T, where S*_*t*_ *is the set of t-specific strings*.

Proof. For the sake of simplicity, assume that *T* = {*t*_1_, *t*_2_}. Let *S* 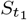 be the set of *t*_1_-specific strings obtained as a solution of the SFS problem on instance ({*t*_1_}, *R*) and similarly let 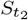 be the set of *t*_2_-specific strings on instance ({*t*_2_}, *R*). We need to prove that given *S*_*T*_ the solution of the SFS problem on instance ({*t*_1_, *t*_2_}, *R*), then 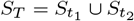. Let us first observe that 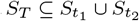 as indeed each string *s* in *S*_*T*_ must be a substring of *t*_1_ or of *t*_2_ and thus *s* is a *t*_1_-specific string or is a *t*_2_-specific string. Hence let us now prove that 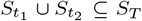. By construction, any *t*_1_-specific string (as well as any *t*_2_-specific string) is a substring of a string in *T* (condition 1) and it is not a substring of any string in *R* (condition 2). Moreover, strings in 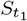 (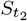, respectively) are substring-free in the sense that each string is not a substring of another one in the same set. We have to prove that any *t*_1_-specific string *x* cannot be a substring of any *t*_2_-specific string *y*, and vice versa (substring-free property). We will prove this by contradiction. Let us assume that *x* is a substring of *y*. By definition *y* is not a substring of *R* which implies that *x* is a substring of *R*: indeed *y* being substring-free, it holds that any substring of *y* is a substring of *R*. But *x* being a *t*_2_-specific string, we obtain a contradiction. At this point, the vice versa is trivial to prove.

## 3 Preliminary concepts

The FMD-index [25] is a data structure based on the FM-index [16] which indexes a set of strings and their reverse complements at the same time, allowing to perform search operations on the index. Differently from the bidirectional BWT [23] which builds two FM-indices, the FMD-index builds a single FM-index for both strands. The FM-index of the collection {*r*_1_, …, *r*_*n*_} of strings of sample *R* is essentially made of the BWT (Burrows Wheeler Transform) of *R* which is itself a permutation *B* of the symbols of *R* obtained from the Generalized Suffix Array (GSA) *SA* of *R*. Indeed, recalling that *SA*[*i*] is equal to (*k, j*) if and only if the *k*-suffix of string *r*_*j*_ is the *i*-th smallest element in the lexicographic ordered set of all suffixes of the strings in *R*, then *B*[*i*] = *r*_*j*_ [|*r*_*j*_ | − *k*], if *SA*[*i*] = (*k, j*) and *k <* |*r*_*j*_ |, or *B*[*i*] = $ otherwise. Given a string *Q*, all suffixes that have *Q* as a prefix appear consecutively in GSA, where they induce an interval [*b, e*) which is called *Q-interval*. Note that the difference *e* − *b*, also called the width of the *Q*-interval is equal to the number of occurrences of *Q* as a substring of some string *r* ∈ *R*. The backward extension operation of an arbitrary character *σ* applied to the *Q*-interval of a string *Q* allows to determine the *σQ*-interval in the index. In particular, iteratively performing the backward operation on a pattern by searching the pattern backwards from its last symbol to its first symbol, allows to find all occurrences of the pattern inside the strings of the reference sample *R* in linear time in the size of the pattern. The FMD-index also allows to apply a forward extension operation of an arbitrary character *σ* to a *Q*-interval of a string *Q* to determine the *Qσ*-interval in the index. The implementation of both forward and backward operations in the FMD-index is obtained by constructing a FM-index for the collection *R* concatenated with the reverse-complement of each string.

In the following, we will simultaneously apply backward and forward extensions on both orientations of a collection of strings. By adopting the same notations as in [25], we keep a triple [*i, j, l*] (called *bi-interval*) that encodes for the *Q*-interval [*i, i* + *l*] and the 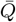-interval [*j, j* + *l*], where 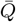 is the reverse complement of string *Q*. Whenever *l* = 0 the *Q*-interval (respectively 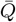-interval) is empty and string *Q* (respectively 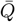) does not occur in *I*_*R*_. We will use notation *t*[*b* : *e*] to denote an interval on string *t*, i.e., *t*[*b* : *e*] is a substring of t, whereas [*i*_*b*_, *j*_*b*_, *l*_*b*_] to denote the corresponding *t*[*b* : *e*]-interval on the index *I*_*R*_.

## 4 Algorithm for sample-specific string detection

We present a novel algorithm (Algorithm 1, Ping-Pong search) to solve the SFS problem between a set of reference strings and a single target string *t* ∈ *T*. Our algorithm computes substring-free *t*-specific strings with respect to the reference sample *R* using the FMD-index of *R*, from now on denoted *I*_*R*_. Note that based on Property 1, it is straightforward to extend the proposed algorithm to solve the SFS problem between a set of reference strings and a set of target strings (i.e., *T*). We will give more details on this at the end of this section.

The following main property which is a direct consequence of the substring-free property of specific strings is used to define the generic iteration step of Algorithm 1.

### Algorithm 1 Computing *t*-specific strings from FMD-index *I*_*R*_

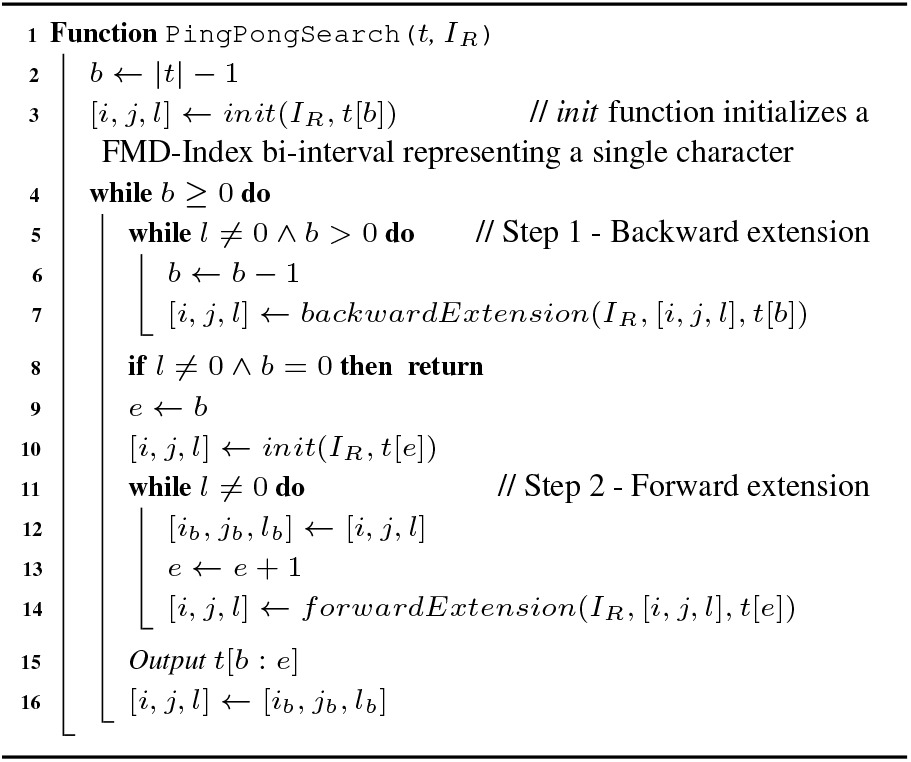

### Lemma 1

*Let R be a collection of strings with FMD-index I*_*R*_ *and let t be a string that does not exist in R. Let x be the rightmost t-specific string currently found in t, where x* = *t*[*b*_*x*_ : *e*_*x*_]. *It must then be the case that any other t-specific string must begin before b*_*x*_. *Assume such a specific string y exists and starts at b*_*y*_, *it must then be the case that y is the shortest prefix of t*[*b*_*y*_ : *e*_*x*_ − 1] *that does not occur in the index*.

Proof. By definition, two specific-strings cannot start at the same position as one cannot be a substring of the other. Thus given *x* the rightmost occurrence of a specific-string in a string, then a second rightmost occurrence *y* of a specific string must start to the left of *b*_*x*_, i.e., given *y* = *t*[*b*_*y*_ : *e*_*y*_] it holds that *b*_*y*_ *< b*_*x*_. By the substring-free property it must be that *t*[*b*_*y*_ : *e*_*x*_] does not occur in the index as it contains substring *x* which does not occur in the index. On the other hand it must be that *e*_*y*_ *< e*_*x*_ otherwise *y* includes *x* as *b*_*y*_ *< b*_*x*_ which is not possible by definition of *x* and *y* as being both specific-strings. Thus *e*_*y*_ *< e*_*x*_ which implies that *y* is a prefix of *t*[*b*_*y*_ : *e*_*x*_ − 1]. Now, it must be the shortest prefix not in the index, otherwise it includes another specific string contradicting the substring-free property, which clearly implies that also *t*[*b*_*y*_ : *e*_*x*_ − 1] is not in the index, thus concluding the proof of the Lemma.

Based on the previous Lemma, given the interval [*b*_*x*_ : *e*_*x*_] of the last detected specific-string, the algorithm will start looking for a new occurrence of a specific-string from the end position *e*_*x*_ − 1.

More precisely, the algorithm keeps track of two search positions *b* and *e* inside *t* which respectively represent the start and end of a substring of *t* that may or may not exist in *I*_*R*_. The algorithm uses the constant-time forward and backward extension operations defined on the FMD-index [25]. Given the index *I*_*R*_ and a triple [*i, j, l*] encoding a *Q*-interval and 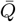-interval, the algorithm alternates between extending the *Q*-interval backward (step 1, lines 5 to 7) and forward (step 2, lines from 11 to 14) to find t-specific strings. Figure 1b illustrates how the algorithm iterates over an input string *t*.

During each iteration of step 1, the algorithm backward extends the *t*[*b* : *e*]-interval of *I*_*R*_ with *t*[*b* − 1] until the backward extension in the index *I*_*R*_ with *t*[*b* − 1] is not possible. In other words, this is equivalent to finding the left maximal match ending at position *e* and extending it one base on the left. Now *t*[*b* − 1 : *e*] is a substring of *t* that is specific to *t*. However, such a substring is not necessarily the shortest, since one of its prefixes may also be specific.

Step 2 initializes *e* to *b* − 1 and then keeps incrementing *e* by one position at a time, and performs a forward extension in *I*_*R*_ for the prefix *t*[*b* − 1 : *e*] for each increment. If the forward extension with *t*[*e* + 1] is not possible in *I*_*R*_, the algorithm stops and returns *t*[*b* − 1 : *e* + 1] as the shortest string beginning from position *b* − 1 that’s not in *I*_*R*_. In other words, we are looking for the longest right maximal match starting at position *b* − 1 and then we are extending it one base on the right. We note that Algorithm 1 outputs substring *t*[*b* : *e*] since *b* (resp. *e*) has been already decremented (resp. incremented) previously in the corresponding `while` (i.e., step 1 for *b* and step 2 for *e*). Finally, since substring *t*[*b* − 1 : *e*] is not *t*-specific and is in the index, it could be extended to the left to compute a new *t*-specific which will eventually overlap the last computed *t*-specific *t*[*b* − 1 : *e* + 1]. Line 16 initializes this process. Observe that Algorithm 1 when processing a string *t* may compute the same specific-string multiple times, even though the output is a set of *t*-specific strings.

### Theorem 1

*Algorithm 1 solves the SFS problem for a string t w*.*r*.*t. a reference set R in time* 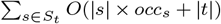, *where occ*_*s*_ *is the number of times a string s is output by Algorithm 1 when processing t*.

Proof. We start by proving correctness and then time complexity. Based on Lemma 1, the Algorithm searches for a new specific-string starting from the end position *e*_*x*_ of the last detected specific-string *x*. The correctness relies on the fact that Algorithm 1 visits from right to left each position *b* of the prefix of length *e*_*x*_ of the input string *t* maintaining the following invariant property: the Algorithm outputs the shortest prefix *t*[*b* : *e*] of *t*[*b* : *e*_*x*_ − 1] which does not occur in the index *I*_*R*_ and *t*[*b* : *e*_*x*_ − 1] is the rightmost string detected as not occurring in the index ending in *e*_*x*_ − 1, if such string exists. Based on Lemma 1, this invariant property allows us to state that the Algorithm for any position *b* outputs the *t*-specific string starting at that position (which is unique by the substring-free property) if any; since all positions of the input string are processed by the Algorithm, at the end it outputs all possible specific-strings. We now show the invariant by analyzing a single iteration. Assume that *b* is a position such that *t*[*b* : *e*] is a *t*-specific string computed when the algorithm visits such a position of *t*. Initially, *b* is the position before the end of *t* (line 2 of the Algorithm). Now, let *k* be the smallest integer (with *k < b*) such that *t*[*b* − *k* : *e* − 1] is the next string *x* not in the index. This is easily detected by backward extension, i.e., by iterating *k* times the loop from line 5 to 7 of the Algorithm. After finding *k*, the algorithm sets *k* ′ = 0 and computes whether *t*[*b* − *k* : *b* − *k* + *k* ′] is in the index for increasing values of *k* ′ and stops as soon as *t*[*b* − *k* : *b* − *k* + *k*] is not in the index thus computing the shortest prefix of *t*[*b* − *k* : *e* − 1] not in the index. This is done by first initializing the *Q*-interval to be the interval for *Q* = *t*[*b*], where *b* = *b* − *k* and then by iterating *k* times the loop from line 11 to line 14 and forward extending the interval. This concludes the proof of the invariant. To prove Algorithm 1 time complexity, observe that it performs a number of backward extensions which is equal to the length of the string *t*, while it performs a number of forward extensions that is *O*(*l*_*b*_) for *l*_*b*_ the length of the specific string given in output when the algorithm is processing the suffix from position *b* of *t*. Thus the time complexity easily follows from the above observation.

### Relaxed Ping-pong Search: a faster heuristic search algorithm

Observe that by Theorem 1 the worst case time required to solve the SFS problem on a single string *t* is *O*(*n*^2^) for *n* being the length of the string *t*, assuming that the index *I* is already available. Note that in the formula 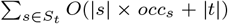, |*s*| can be *O*(*n*) in the worst case and 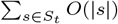 can achieve the bound of *O*(*n*^2^) since the strings in *S*_*t*_ span positions of the string *t* that are overlapping and we can have *O*(*n*) strings in *S*_*t*_ each of length *O*(*n*). See Supplementary Section 1.1 for an example. This clearly implies a quadratic time for solving the SFS problem when the input is no longer a single string *t* but a collection *T* of strings of total length *n*.

We consider a simple variant of Algorithm 1 that leads to a linear-time complexity by avoiding the computation of specific strings that occur in overlapping positions of the original string *t*. The variation is simply obtained from the pseudo-code of Algorithm 1 by deleting instruction 12 and replacing line 16 with the instruction [*i, j, l*] ←*init*(*I*_*R*_, *t*[*b* − 1]). This implies that the search procedure of *t*-specific strings starts from the left of the beginning of the last detected one in *t*, instead from position *e*. We call this procedure the *relaxed Ping-pong Search*.

It is easy to verify that the relaxed version of Algorithm 1 is linear in the size of string *t*. Indeed, in the worst case it performs two index queries per symbol of the input string: each character is searched in the index one time during the backward extension and one time during the forward extension (see Figure 1b). Formally, when estimating the formula 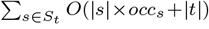 of Theorem 1 in this variant, strings in *S*_*t*_ occur in positions of *t* that are disjoint and thus in the worst case the sum of the sizes of strings in *S*_*t*_ is 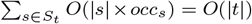, thus proving that the time complexity of the algorithm is linear in the size of the input string.

*Relaxed output set upper-bounded by the edit distance* The edit-distance is a well known measure in the comparison of two genome sequences. By counting the minimum number of nucleotide insertions, deletions and changes that transform a genome *t* into *r*, the edit distance between *t* and *r*, denoted by *D*(*t, r*) is clearly an upper bound for the number of positions with variations in *t* w.r.t. to *r*. In the following we show that for a pair of strings *t* and *r*, each *t*-specific string returned by the relaxed version of Algorithm 1 corresponds to at least one edit operation that changes *t* into *r*, thus showing that *D*(*t, r*) is an upper bound on the size of its output set. Observe that the relaxed version of Algorithm 1 computes a subset of the *T*-specific strings w.r.t. *R* that has the substring-free property.

#### Theorem 2

*Given two strings t and r*, |*S*_*t*_| *the size of the set of strings S*_*t*_ *returned by the relaxed Ping-pong search with respect to r, then* |*S*_*t*_| ≤ *D*(*t, r*).

Proof. Since the set *S*_*t*_ consists of string induced by non-overlapping intervals of sequence *t*, any edit operation changes |*S*_*t*_| by at most 1. The minimum set of edit operations to convert *t* to *r* (i.e., *D*(*t, r*)) will transform *t* = *t*_0_ into successive strings *t*_1_, *t*_2_, …, *t*_*D*(*t,r*)_ and eventually *t*_*D*(*t,r*)_= *r*. For each operation, the successive sets of relaxed Ping-pong strings for *t*_2_, …change in cardinality by at most 1, i.e. 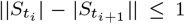 for 0 ≤ *i < D*(*t, r*). Observe that *D*(*t, r*) = 0 implies 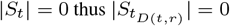. Thus, total size of |*S*_*t*_| could not have been more than *D*(*t, r*) to start.

### Implementation details

We implemented Algorithm 1 in `C++` based on code from `ropeBWT2` [27]. After creating the index of the reference set *R*, our code executes Algorithm 1 on each target string *t* ∈ *T* while also keeping track of the number of times each specific string is seen (Figure 2). As each target string can be processed independently, our code is embarrassingly parallel. Once all target strings have been analyzed, a post-processing step combines the smaller solutions into the final solution of the SFS problem. In order to remove specific strings produced by sequencing errors when our method is run on WGS data, the post-processing step can filter out all the specific strings occurring less than *τ* times, with *τ* being a user-defined cutoff. Our implementation is freely available at https://github.com/Parsoa/PingPong.

**Fig 2:**
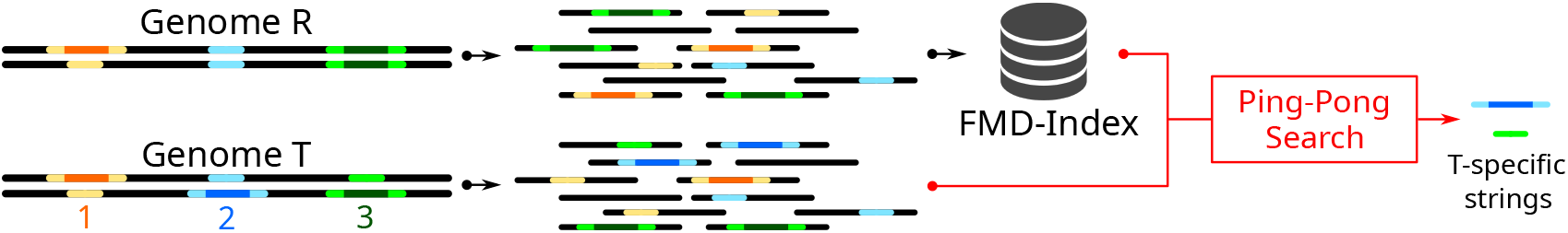
Illustration of sample-specific string detection. Two genomes *R* and *T* are depicted. With respect to genome *R*, site 1 has no variation in *T*, site 2 is an heterozygous insertion in *T*, and site 3 is an heterozygous deletion in *T*. Our pipeline aims to detect *T*-specific strings by (a) indexing the reads sequenced from *R* with a FMD-Index and (b) analyzing the reads sequenced from *T* with our novel *Ping-pong search* algorithm. We note that, for ease of presentation, we depict at the end of the pipeline a single *T*-specific string per site even though multiple *T*-specific strings may actually be reported for each site.

## 5 Results

### 5.1 Specific string detection in simulated human HiFi trio

We used simulations to test the performance of our proposed method in detecting *de novo* SVs in WGS trios (i.e., proband, mother and child). We mutated the GRCh38 genome randomly with 6115 insertions and deletions from the 1KG project [10] to produce two haplotypes for each parent. We limited the simulations to chromosomes 1-5. We then simulated the child genome by inheriting variants from the parents and considering recombination inside each chromosome. Finally, we introduced an additional 17,595 randomly-generated *de novo* structural variants equally divided between insertions, deletions and inversions into the child genome impacting 7,913,593 base-pairs. See Supplementary Figure S1 for the distribution of lengths of simulated SVs.

We simulated reads from the father, mother and child genomes at different coverage levels (5x, 10x, 20x and 30x) for each haplotype using `PBSIM` [36] with sequencing error rate and read length distribution similar to real HiFi data. Specifically, we sampled these parameters from the HGSVC2 PacBio HiFi reads for the HG00733 sample [40] with the error rate averaging at 0.1%. All three samples were error corrected using `ntEdit` [48] to remove sequencing errors. The combined reads of the father and mother were indexed using FMD-index and we searched for child-specific strings using Algorithm 1 (exact version).

We measured the accuracy of the method using two metrics of **recall** and **precision**. Recall is defined as the percentage of *de novo* variants that are covered with child-specific strings and precision is defined as the percentage of child-specific strings that cover a *de novo* variant. We test the performance of the method for different *τ* cutoff values (2 ≤ *τ* ≤ 6) to study the relationship between this parameter and sequencing coverage levels and to measure our method’s sensitivity (Figure 3). While the high coverage simulations (30x, 20x and 10x) have constantly high recall rates regardless of *τ*, the low-coverage 5x sample’s recall drops significantly with larger cutoff values.

**Fig 3:**
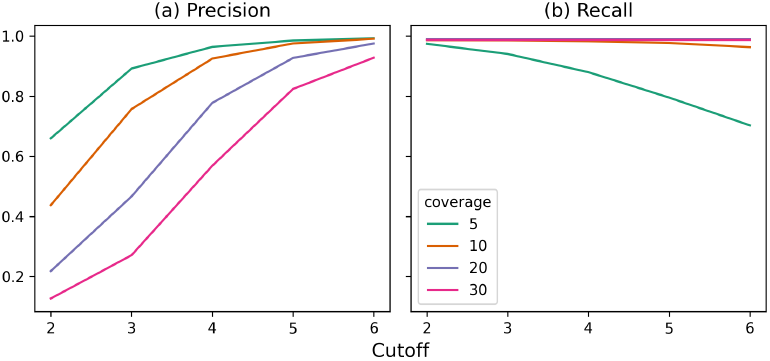
Precision (a) and recall (b) calculated for different coverage levels (5x, 10x, 20x and 30x per haplotype) and cutoff values 2 ≤ *τ* ≤ 6 in simulation.

We analyzed the child-specific strings from the 30x simulation using *τ* = 5 in more detail. A total of 14 381 350 child-specific strings were retrieved with 2 052 144 remaining after filtering low-abundance strings. The selected child-specific strings achieved *>* 98% recall and 82% precision at recovering simulated *de novo* structural variations. To better demonstrate the usefulness of child-specific strings, we compared the performance of the strings generated using both the exact and relaxed versions of Algorithm 1 on the 30x simulation against child-specific *k*-mers of fixed sizes 32bp and 101bp with abundance of at least 5. Child specific *k*-mers were calculated using `KMC3` [22] by subtracting the set of parent *k*-mers from the set of child *k*-mers. We calculated precision and recall by mapping the *k*-mers and SFS strings to the child haplotypes with `BBMap` [6]. We observe that SFS consistently performs better than fixed-length *k*-mers. The results can be seen in Table 1.

**Table 1.** Comparison of performance of SFS and fixed-length *k*-mers in the 30x simulation with *τ* = 5.

| Method | Sequences |  |  | Variants |  |
| --- | --- | --- | --- | --- | --- |
|  | Total | Covering | Precision | Covered | Recall |
| Alg. 1 (exact) | 1 690 675 | 2 052 144 | 82.38% | 17367 | **98.70%** |
| Alg. 1 (relax) | 377 007 | 366 390 | **97.18%** | 17365 | 98.69% |
| 32-mers | 5 089 147 | 3 053 969 | 59.91% | 17317 | 98.42% |
| 101-mers | 7 060 167 | 3 453 211 | 48.91% | 17348 | 98.59% |

We further analyzed the qualities of the alignments of child-specific strings against all three genomes in the trio (Supplementary Figure S2). Alignment quality is evaluated based on the number of bases that do not match. More than 83% of child-specific strings map perfectly to the child genome, and no (zero) string has a mismatch-free mapping to either parent genomes, indicating that the strings are truly child-specific.

Finally, we re-ran the simulation at 30x coverage without incorporating any sequencing errors in the trio. In this scenario, the simulated SVs are the sole source of novel sequences in the child compared to the parents and therefore we expect every recovered SFS to cover a variant. Analyzing the 1 720 395 child-specific strings retrieved in this scenario indeed yields a precision of 100.0%. However, the recall remains the same as in the case with sequencing errors, at 98.70%. This is because some variants don’t produce novel sequences and thus cannot be captured with our approach.

### 5.2 Specific string detection in real human HiFi data

We performed an extensive evaluation of sample-specific strings using real HiFi data to assess their ability to compare two individuals of different populations. We considered the HG00733 child (Puerto Rican trio) and the NA19240 child (Yoruba trio). For both these individuals, the HGSVC2 [40] provides a PacBio HiFi 30x sample. Supplemental Figure S3 reports the length distribution of the considered samples.

After correcting both samples with `ntEdit` [48], we indexed the NA19240 sample and we searched for HG00733-specific strings (from now on we will refer to these strings simply as ‘specific’) using both the exact and the relaxed version of our algorithm. Supplemental Table S1 reports the running times and the peak memory usage of our pipeline; the creation of the FMD-Index was the most time-consuming step. Based on the results on simulated data (Figure 3) and the coverage of the two samples (30x), we considered all specific strings occurring more than 5 times. The main goal of this postfiltering is to remove from downstream analyses specific strings that with high probability are the result of sequencing errors. Using the exact (relaxed, respectively) version of our algorithm we retrieved 34 219 149 (7 125 436, respectively) strings. Supplemental Figure S4 reports information on the lengths and the abundances of these strings. As expected, the exact version of our algorithm is slower and retrieves more strings than the relaxed one.

#### Contigs-based analysis

We first analyzed the quality of HG00733-specific strings by checking whether they are effectively specific to the HG00733 child. To do so, we aligned the strings to the contigs provided by the HGSVC2 consortium of the two individuals and we counted base differences (substitutions, insertions, deletions, and clips) within alignments. We mapped strings shorter than 500bp with `BBMap` [6] and longer ones with `minimap2` [6]. We used two different aligners since `BBMap` showed higher sensitivity in mapping short (*<* 50bp) strings. Figure 4 (a/b) shows the results of this analysis for the exact version of our algorithm (see Supplemental Figure S5 for the relaxed results). A total of 33 964 009 specific strings were mapped to the HG00733 contigs and 27 326 747 (80%) of these were aligned perfectly, i.e., without any base difference. On the other hand, 33 932 307 specific strings were mapped to the NA19240 contigs but only 158 094 (0.4%) of these were aligned perfectly.

**Fig 4:**
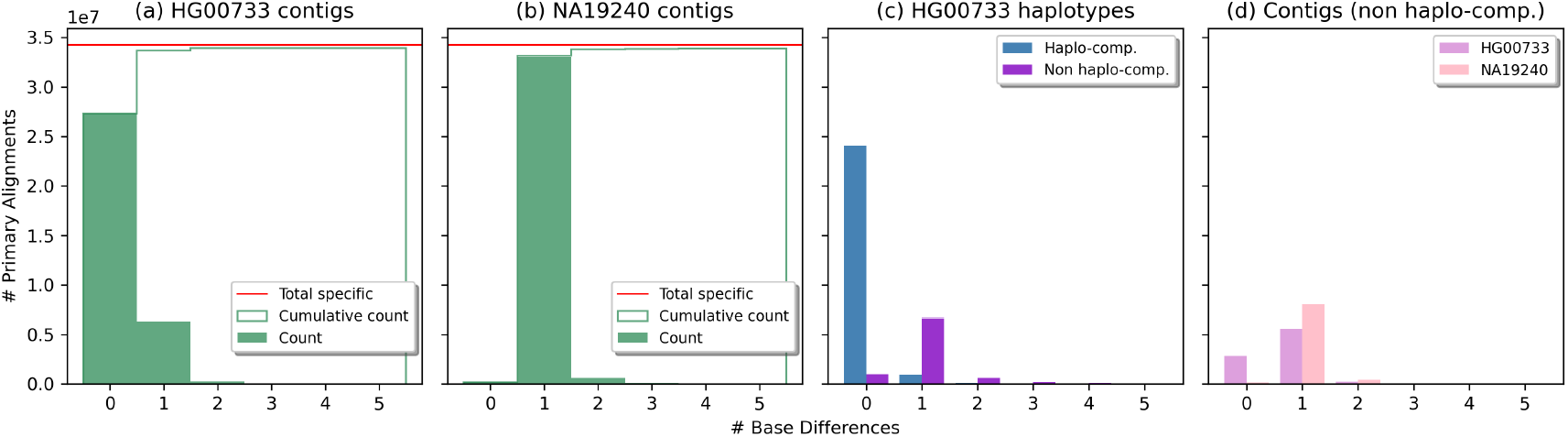
Results on exact HG00733-specific strings. Panels (a, b): comparison of the quality of specific strings alignments computed against the HG00733 contigs (a) and the NA19240 contigs (b). Panel (c): comparison of the qualities of the specific string alignments representing (Haplo-compatible) and not representing (Non haplo-compatible) a specific portion of HG00733 haplotypes. Panel (d): comparison of the qualities of non haplo-compatible specific string alignments computed against the HG00733 contigs and the NA19240 contigs. Quality is expressed as number of base differences (mismatches, insertions, deletions, and clips).

To summarize the results of this contigs-based analysis, we introduced the **C-precision** (contigs-based precision) metric. Based on the alignments to the contigs, it computes the fraction of HG00733-specific strings that align perfectly to HG00733 contigs and not perfectly to NA19240 contigs. Out of 27 326 747 specific strings aligned perfectly to HG00733 contigs, 132 031 aligned perfectly also to NA19240 contigs. The exact version of our algorithm therefore achieved a C-precision of 79.47%. On the other hand, the relaxed version achieved a C-precision of 90.61%. This was expected since the relaxed version of our algorithm retrieves a lower number of specific strings easily achieving a higher precision at the expense of, as we will see in the next section, a lower recall (see Table 2). This analysis shows that the strings output by our algorithm are effectively specific to the HG00733 and may be effectively used to characterize differences between the two individuals.

**Table 2.**
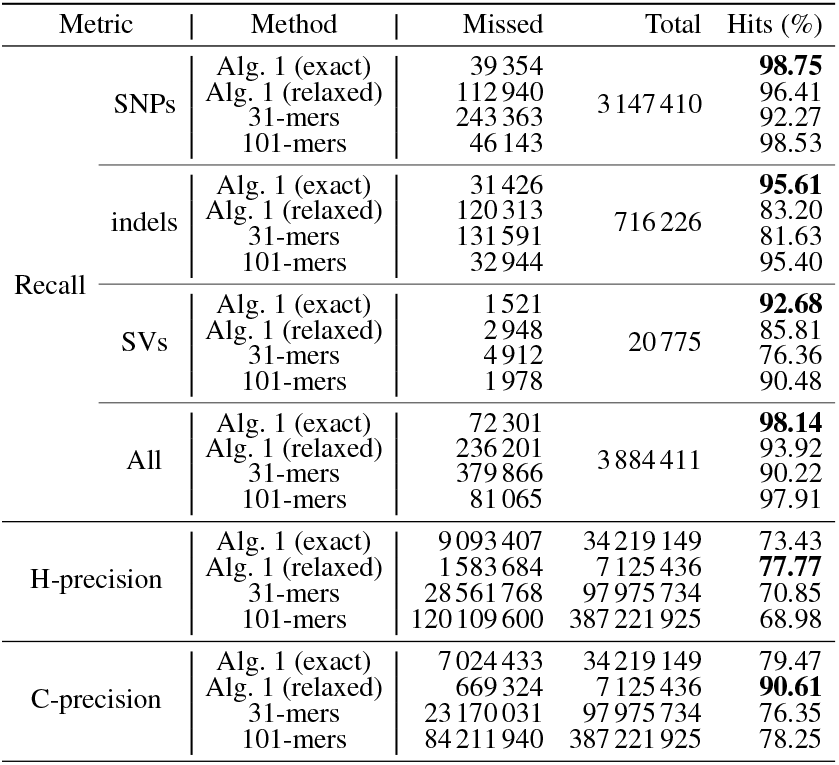
Variant analysis on real human HiFi data. Recall is the fraction of known alleles specific to HG00733 (w.r.t. NA19240) overlapped by at least one HG00733-specific string (or specific *k*-mer). For the sake of completeness, we reported the recall values for alleles coming from SNPs, indels (2-49bp), and SVs (≥ 50bp), as well as all considered specific alleles. H-precision (Haplotype-aware precision) is the fraction of HG00733-specific strings (or HG00733-specific *k*-mers) representing a portion of its haplotypes that is specific w.r.t. the NA19240 haplotypes. C-precision (Contig-based precision) is the fraction of HG00733-specific strings (or *k*-mers) aligning perfectly only to HG00733 contigs (and with errors to NA19240 contigs).

#### Haplotypes-based analysis

We evaluated the effectiveness of HG00733-specific strings in covering variant alleles that are specific to the considered individual. To do so, we considered the phased callset provided by the HGSVC2 consortium [15] and, after filtering out overlapping variations, we extracted for each variation and for each haplotype the set of alleles that are present in the HG00733 child but not in the NA19240 (we will refer to these alleles as *HG00733-specific* or simply *specific alleles*). Therefore, each variation may have 0, 1 or 2 specific alleles. For instance, if a variation has genotype 0|2 in the HG00733 child and 1|1 in the other child, we considered alleles 0 and 2 as specific to the HG00733. Table 2 (column *Total*) reports the number of specific alleles we considered in our analysis. We classified each allele with respect to the type of its originating variant (following the classification in [15]): SNPs, indels (insertions and deletions of 1-49 bp), and SVs (insertions and deletions of ≥ 50bp), which include copy number variants and balanced inversion polymorphisms.

Considering the entire set of known variations, we built the haplotypes of the HG00733 individual using `BCFtools` and then we aligned the HG00733-specific strings (occurring more than 5 times) to them using `BBMap` (strings ≤ 500bp) and `minimap2` (strings *>* 500bp). Finally, we used `BEDtools` [42] (*intersect* sub-command) to find the overlaps between the alignments and the considered alleles.

We evaluated the quality of our specific strings in terms of **recall**, i.e., number of specific alleles effectively intersected by at least one alignment, and **H-precision** (Haplotype-compatible precision), i.e., the number of specific strings representing a specific portion of a haplotype of the HG00733 child. By “specific portion” we mean a subsequence of a HG00733 haplotype induced by a set of variations that is different from the subsequence of any NA19240 haplotype induced by the same set. Table 2 reports the results of this analysis. We introduced the H-precision measure since close alleles (especially SNPs) on a haplotype of one individual may result in a specific string even when neither alleles are specific. Indeed, a set of close alleles may be shared between two individuals but in one individual they may be on the same haplotype whereas in the other one on different haplotypes. Consider for instance two nearby variants with genotypes 0|1 and 0|1 in one individual and 1|0 and 0|0 in the other. In this case, the haplotype containing alleles 1 and 1 is specific to the first individual even though single alleles are not.

Remarkably, the set of specific strings computed by our method (exact version) intersect most of the HG00733-specific alleles (*>* 98%), covering nearly all alleles coming from SNPs and indels (*>* 98% and *>* 95%, respectively) and most of alleles coming from SVs (*>* 92%). We observed that a majority of the variants not covered by the sample-specific strings were indels in stretches of A or T sequences, likely addressable through improvements in homopolymer error correction.

Out of the 34 219 149 specific strings retrieved by the exact version, 73.43% of them represent a specific portion of the HG00733 haplotypes (H-precision). Figure 4 (c) reports the comparison in terms of base differences between the alignments representing specific portions of the haplotypes (denoted as “haplo-compatible”) and those that do not (denoted as “non haplo-compatible”). As expected, the vast majority of the haplo-compatible strings are aligned perfectly to the haplotypes whereas the vast majority of non haplo-compatible strings are aligned with errors.

To better investigate why ∼ 27% of the specific strings align well to the HG00733 haplotypes but do not represent a specific portion of them (accordingly to the considered VCF), we aligned those strings to the contigs of the two individuals. Figure 4 (d) reports the results of this analysis. 2 885 356 strings were aligned perfectly to the HG00733 contigs whereas only 208 330 were mapped perfectly to the NA19240. Moreover, ∼ 1.8 million specific strings align perfectly to the HG00733 contigs but not to its haplotypes. This leads us to conjecture that a portion of those strings correspond to true variants missing from the VCF.

Results on strings retrieved by the relaxed algorithm follow the same trend (see Supplemental Figure S5). They however achieve higher H-precision and lower recall than the exact version (see Table 2), likely due to a lower number of strings returned. Moreover, the relaxed algorithm may fail in covering close variations: if two variations are too close to each other, strings retrieved by our relaxed algorithm may cover only the right-most variation (due to its right-to-left traverse of input strings). See Supplemental Figure S6 for an example.

To put our results in perspective, we compared them with a *k*-mer method. Similarly to a HG00733-specific string, a *HG00733-specific k-mer* is a *k*-mer occurring in the HG00733 sample and not in the NA19240. To compute the set of specific *k*-mers we first counted all *k*-mers occurring more than 5 times in the two samples independently with `KMC3` [22] and then we retrieved the *k*-mers present only in the HG00733 sample by subtracting the two sets (`kmc_tools` *kmers_subtract* operation). A total of 97 975 734 HG00733-specific *k*-mers (*k* = 31) were retrieved. We then mapped those to HG00733 haplotypes with `BBMap` and evaluated their recall and H-precision similarly to HG00733-specific strings. Table 2 reports the results of this analysis. HG00733-specific 31-mers achieved lower recall and H-precision than HG00733-specific strings, although their computation is faster (8h for *k*-mers vs 28–37h for Ping-Pong, see Supplemental Table S1). The poor performance of 31-mers can be explained by their length: a 31-mer located at a variant position might occur elsewhere in the genome, whereas a longer string would be unique. We note that long (>500bp) HG00733-specific strings retrieved by our (exact) algorithm cover ∼ 1.5% of indels and SVs not covered by shorter ones, proving that in some scenarios longer strings are needed to effectively cover a variation.

For this reason, we also performed an analysis using longer *k*-mers (*k* = 101). A total of 387 221 925 101-mers were retrieved. However `BBMap` failed to align that many *k*-mers in reasonable time. We therefore aligned them with `BWA-MEM` [26] and computed their recall and H-precision. Results of this analysis can be found in Table 2. Thanks to their length, 101-mers are able to cover more variations than 31-mers but not as many as our (exact) specific strings which are of variable length, sometimes longer than 101bp. Moreover, 101-mers are less precise than (exact) HG00733-specific strings: indeed, due to their overlapping nature, a false variant (e.g., a sequencing error) will in the worst case yield 101 false specific 101-mers. We therefore mapped the specific *k*-mers to the contigs of the two individuals and we computed their C-precision (fraction of specific *k*-mers mapping perfectly only to HG00733 contigs). Similarly to specific strings, C-precision of 31-mers and 101-mers is higher then their H-precision (see Table 2), proving one more time that the considered VCF may be incomplete.

Finally, in an attempt to reduce the number of strings obtained using the *k*-mer method, we assembled the 31-mers and the 101-mers into unitigs (which correspond to maximally extending *k*-mers using their (*k* − 1)-overlaps and stopping at any variation) using `BCALM2` [11] and we computed their recall and H-precision. Results of this analysis, can be found in Supplemental Table S2. Surprisingly, assembling the *k*-mers into unitigs did not improve their overall accuracy.

## 6 Discussion

We have presented a novel approach using the sequence differences between multiple samples, denoted as substring-free samples-specific (SFS) strings, for performing comparative genome analysis between two or more whole-genome sequenced samples. We have shown that these SFS strings are a comprehensive representation of variants between samples of interest. In practice the proposed approach is capable of finding sequences that match the breakpoints of most variants.

We have proposed SFS strings as a novel theoretical concept to be utilized in comparing two or more sequences to find differences between them. We also developed novel methods for efficient finding of SFS strings denoted as Ping-Pong algorithm. The SFS strings can have a wide range of applicability in comparative genome analysis. We have shown the applicability of SFS strings in two genomics applications. First, SFS are shown to be capable of discovery of *de novo* variants in a child genome versus parents (see Section 5.1). Second, SFS are shown to be capable of discovery of variants between two WGS samples (see Section 5.2). We are also developing approaches to utilize SFS strings to accurately predict structural variants in WGS samples that are not found using traditional read mapping based approaches.

The proposed approach improves upon using fixed length sequences (i.e., *k*-mers) for comparative genome analysis in three aspects: 1) recall, 2) precision and 3) number of returned strings. 1) SFS sequences cover a higher fraction of true difference between two genomes than constant length *k*-mers (i.e., higher recall). Mainly, this is due to their variable-length nature which increases our power in finding strings representative of differences between genomes in repetitive regions (e.g., segmental duplications). 2) Furthermore, our experiments have also indicated that SFS sequences have a higher precision than fixed length *k*-mers (k = 31 or 101). 3) Finally, our exact algorithm returned between 3x-10x less strings than with *k*-mers, making results more amenable to further analysis. For instance, we could not exhaustively map the results of the 101-mer analysis in reasonable time (*<* 1 week).

The proposed approach also has several major advantages over traditional mapping based approaches for comparative genome analysis. First, it is not dependent on predicting variants in each sample for comparative genome analysis. Thus, its performance is not impacted by the biases in variant prediction methods. Second, the proposed approach does not utilize mapping of the reads. Hence, ambiguities in read mappings or biases in mapping algorithms will not impact the results of the proposed method.

One of the main limitations of the proposed method is its reliance on reads with low sequencing error (e.g., HiFi reads). To be able to accurately predict SFS strings from reads with sequencing errors we need to utilize an error correction tool such as `ntEdit`. This method is not expected to translate well to higher error-rate long reads, unless correction yields nearly perfect reads. Furthermore, the possibility of a sample-specific sequence in the target genome being fragmented between multiple reads can result in a potential reduction of overall recall. Note, that there is an interplay between the depth of coverage, lengths of SFS strings, sequencing error rate, cutoff value *τ*, and the recall of the proposed approach. Another shortcoming is the 3x longer running time of the relaxed algorithms compared to *k*-mers.

A potential venue for future research could be to investigate the connection between SFS strings and other related but different concepts in stringology, such as maximal exact/unique matches, minimum unique substrings [50], and shortest uncommon superstrings.

Remarkably the proposed approach is able to recover SFS strings that cover over 98% of the variants between two or more samples. However, we did observe that a majority of the missed variants strings were embedded in homopolymer stretches of A or T that are enriched in potential sequencing errors. Thus, improving the ability of the method to handle sequencing errors would not only extend the reach of our method but also will improve the recall of the method.

## 7 Funding

This project has received funding from the European Union’s Horizon 2020 research and innovation programme under the Marie Skłodowska-Curie grant agreement number [872539], ANR Inception (ANR-16-CONV-0005), and ANR Prairie (ANR-19-P3IA-0001).

## Footnotes

1 https://simpsonlab.github.io/2015/06/15/merging-sv-calls/

## Notes

### Competing Interest Statement

The authors have declared no competing interest.

