## Supplementary Material for "Comparative genome analysis using sample-specific string detection in accurate long reads"

### 1 Additional details on algorithm complexity

#### 1.1 Example of a Ping-pong search in quadratic time

Example 1. *An example of string where the* **Ping-pong Search** *works in quadratic time in the length of the input target $t$ is given by the string $TT\{ACG\}^n$ while the reference $R$ consists of the unique string $TT\{ACG\}^{\frac{n}{2}}$. Observe that $G\{ACG\}^{\frac{n}{2}-1}A$ is a specific-string that occurs at least $\frac{n}{2}-2$-times when the target is processed. Observe that we have multiple occurrences of the same specific-strings and moreover they pairwise overlap. It is interesting to note that in the Relaxed Ping-pong Search each of the specific-string mentioned above have a single occurrence on the input given by the target $t$ and the reference $R$.*

### 2 Additional results and plots for simulated data experiments

#### 2.1 Distribution of simulated SVs

Supplementary Figure S1 shows the distribution of the lengths of simulated SVs. All *de novo* simulated SVs have lengths between 50bp to 1250bp. The simulated SVs were equally divided into the three types of deletion, insertion and inversion.

#### 2.2 Analysis of mapping quality of child-specific strings to parent and child genomes

Figure S2 shows the quality of the alignments of child-specific strings to all three genomes in the trio for the 30x simulation with $\tau = 5$. Alignment quality is calculated based on the number of mismatched bases. As expected, majority ($> 83\%$) of child-specific strings map perfectly to the child genome, while zero strings have a mismatch-free mapping to either parent genomes, indicating that the strings are truly child-specific.

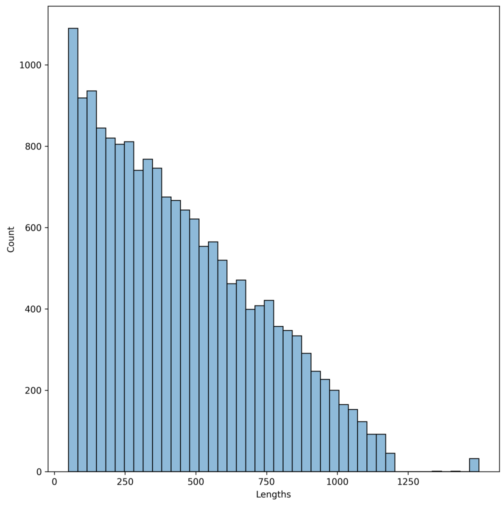


Fig. S1: Distribution of the lengths of simulated SVs.

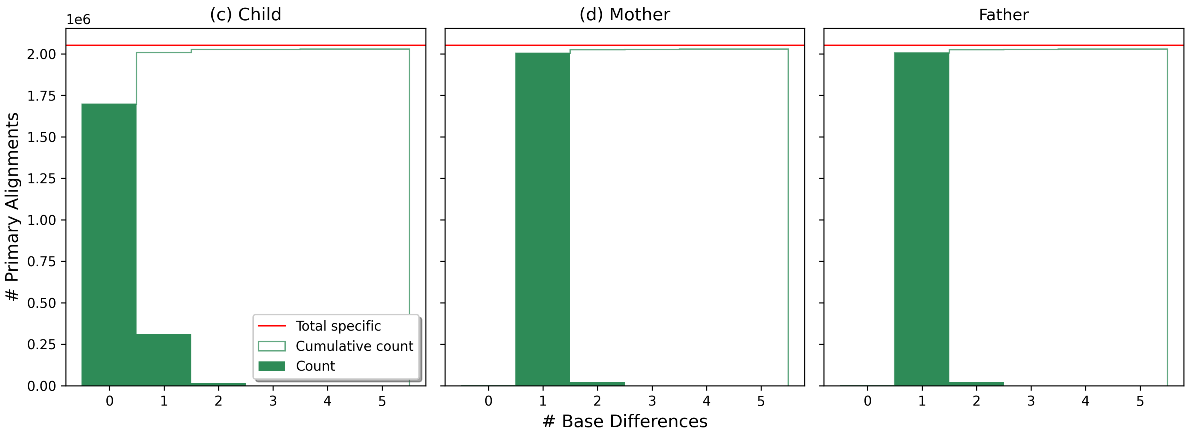


Fig. S2: Comparison of the quality of the alignments of child-specific strings to all three genomes in the trio for the 30x simulation with $\tau = 5$.

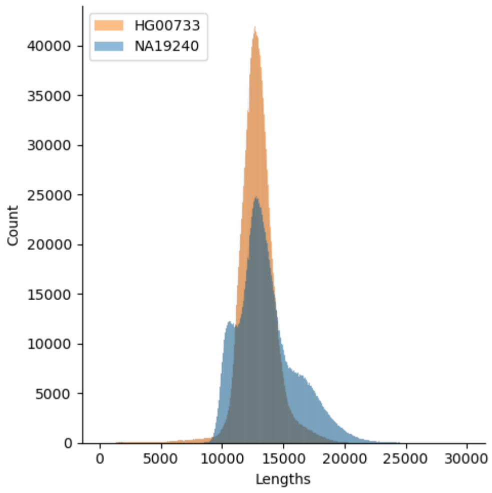


Fig. S3: Read length distribution

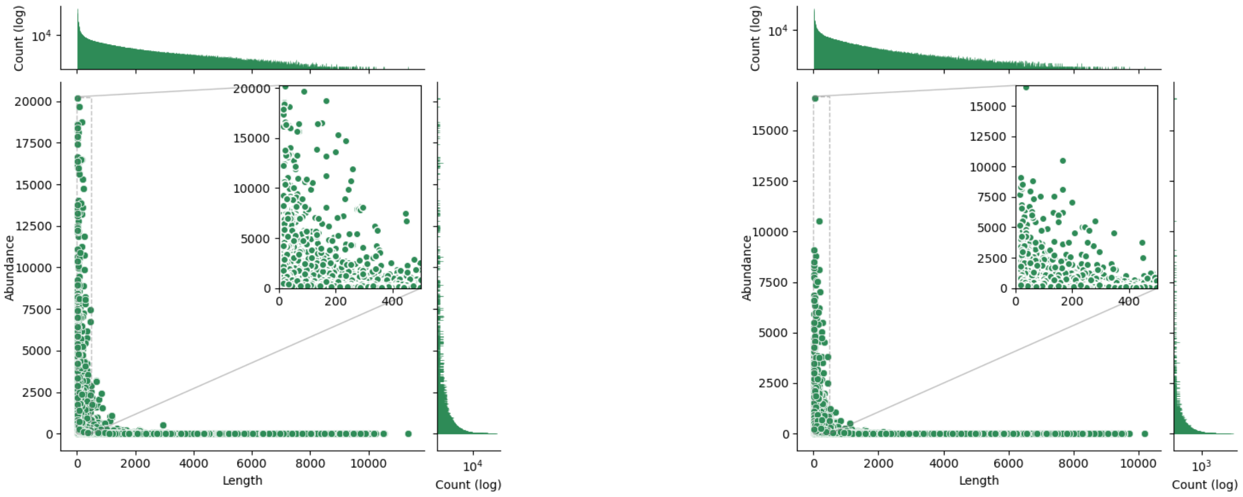


(a) Exact Ping-Pong Search

(b) Relaxed Ping-Pong Search

Fig. S4: Correlation between length and abundance of HG00733-specific strings occurring at least 5 times: 34 219 149 (7 125 436) strings for the Exact (Relaxed) version. For ease of presentation, we zoomed the region concerning strings of length $\leq$ 500bp: 33 600 277 (6 750 216) strings for the Exact (Relaxed) version.

| | | Samples correction | NA19240 indexing | HG00733-specific retrieval | Total Time | Peak RAM |
|---|---|---|---|---|---|---|
| Ping-Pong | Exact | 05:03 | 20:30 | 11:59 | 37:32 | 242 |
| | Relaxed | | | 03:01 | 28:34 | 32 |
| $k$-mers | 31-mers | | - | 03:25 | 08:28 | 12 |
| | 101-mers | | | 03:10 | 08:13 | 24 |

Table S1. Running time (hh:mm) and peak memory (GB) of our pipeline and the $k$-mers pipeline (on real data). We used 16 threads where possible.

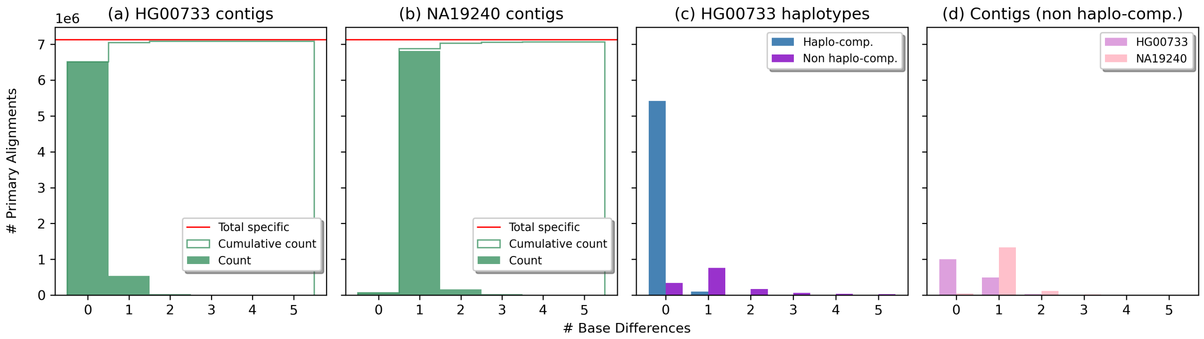


Fig. S5: Results on relaxed specific strings. Panels (a, b): comparison of the quality of HG00733-specific strings alignments computed against the HG00733 contigs (a) and the NA19240 contigs (b). Panel (c): comparison of the qualities of the HG00733-specific string alignments representing (Haplo-compatible) and not representing (Non haplo-compatible) a specific portion of HG00733 haplotypes. Panel (d): comparison of the qualities of non haplo-compatible HG00733-specific string alignments computed against the HG00733 contigs and the NA19240 contigs. Quality is expressed as number of base differences (mismatches, insertions, deletions, and clips).

| Metric | | Method | Missed | Total | Hits (%) |
|---|---|---|---|---|---|
| Recall | SNPs | Alg. 1 (exact) | 39 354 | 3 147 410 | **98.75** |
| | | Alg. 1 (relaxed) | 112 940 | | 96.41 |
| | | 31-mers | 243 363 | | 92.27 |
| | | 101-mers | 46 143 | | 98.53 |
| | | 31-tigs | 267 078 | | 91.51 |
| | | 101-tigs | 55 627 | | 98.23 |
| | indels | Alg. 1 (exact) | 31 426 | 716 226 | **95.61** |
| | | Alg. 1 (relaxed) | 120 313 | | 83.20 |
| | | 31-mers | 131 591 | | 81.63 |
| | | 101-mers | 32 944 | | 95.40 |
| | | 31-tigs | 175 892 | | 75.44 |
| | | 101-tigs | 31 705 | | 95.57 |
| | SVs | Alg. 1 (exact) | 1 521 | 20 775 | **92.68** |
| | | Alg. 1 (relaxed) | 2 948 | | 85.81 |
| | | 31-mers | 4 912 | | 76.36 |
| | | 101-mers | 1 978 | | 90.48 |
| | | 31-tigs | 6 383 | | 69.27 |
| | | 101-tigs | 2 698 | | 87.01 |
| | All | Alg. 1 (exact) | 72 301 | 3 884 411 | **98.14** |
| | | Alg. 1 (relaxed) | 236 201 | | 93.92 |
| | | 31-mers | 379 866 | | 90.22 |
| | | 101-mers | 81 065 | | 97.91 |
| | | 31-tigs | 449 353 | | 88.43 |
| | | 101-tigs | 90 030 | | 97.68 |
| H-precision | | Alg. 1 (exact) | 25 125 742 | 34 219 149 | 73.43 |
| | | Alg. 1 (relaxed) | 5 541 752 | 7 125 436 | **77.77** |
| | | 31-mers | 69 413 966 | 97 975 734 | 70.85 |
| | | 101-mers | 267 112 325 | 387 221 925 | 68.98 |
| | | 31-tigs | 3 075 055 | 5 839 695 | 52.66 |
| | | 101-tigs | 3 114 210 | 5 281 605 | 58.96 |
| C-precision | | Alg. 1 (exact) | 7 024 433 | 34 219 149 | 79.47 |
| | | Alg. 1 (relaxed) | 669 324 | 7 125 436 | **90.61** |
| | | 31-mers | 23 170 031 | 97 975 734 | 76.35 |
| | | 101-mers | 84 211 940 | 387 221 925 | 78.25 |
| | | 31-tigs | 2 563 021 | 5 839 695 | 56.11 |
| | | 101-tigs | - | - | - |

Table S2. Variant analysis on real human HiFi data (complete table). Recall is the fraction of known alleles specific to HG00733 (w.r.t. NA19240) overlapped by at least one HG00733-specific string (or specific $k$-mer or specific unitigs). For the sake of completeness, we reported the recall values for alleles coming from SNPs, indels (2-49bp), and SVs ($\geq 50bp$), as well as all the considered specific alleles. H-precision (Haplotype-aware precision) is the fraction of HG00733-specific strings (or HG00733-specific $k$-mers or HG00733-specific unitigs) representing a portion of its haplotypes that is specific w.r.t. the NA19240 haplotypes. C-precision (Contig-based precision) is the fraction of HG00733-specific strings (or $k$-mers or unitigs) aligning perfectly only to HG00733 contigs (and with errors to NA19240 contigs). C-precision for 101-tigs is not present since `minimap2` crashed while mapping long unitigs to the contigs.

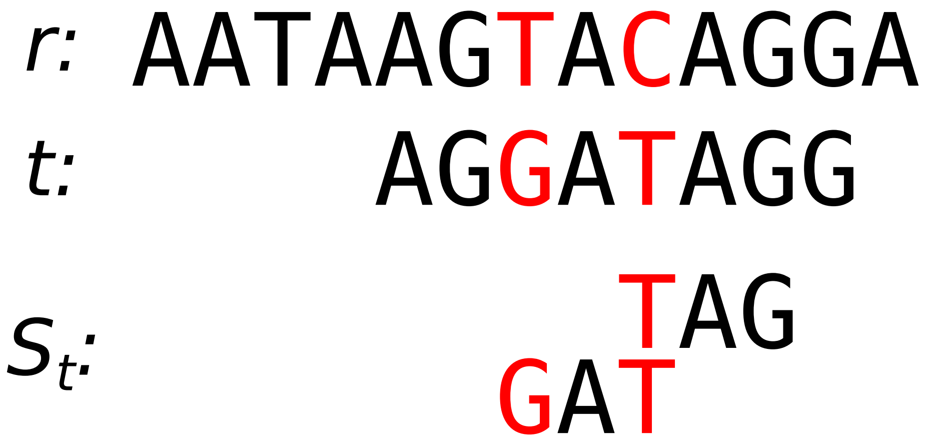


Fig. S6: Let $r$ and $t$ be the reference and the target strings, respectively. We have two variations (SNPs) in $t$. The exact version of our algorithm retrieves the set $S_t$ of $t$-specific strings that cover both variation. On the other hand, the relaxed version would retrieve just the first string since the second one overlaps with the first one. In this case, the set of $t$-specific strings covers only the right-most variation (C>T).
